# Neural responses to heartbeats as a mechanism for distinguishing self from other during imagination

**DOI:** 10.1101/414573

**Authors:** Mariana Babo-Rebelo, Anne Buot, Catherine Tallon-Baudry

## Abstract

Imagination is an internally-generated process, where one can make oneself or other people appear as protagonists of a scene. How does the brain tag the protagonist of an imagined scene, as being oneself or someone else? Crucially, neither external stimuli nor motor feedback are available during imagination to disentangle imagining oneself from imagining someone else. Here, we test the hypothesis that an internal mechanism based on the neural monitoring of heartbeats could distinguish between self and other. 23 participants imagined themselves (from a first-person perspective) or a friend (from a third-person perspective) in various scenarios, while their brain activity was recorded with magnetoencephalography and their cardiac activity was simultaneously monitored. We measured heartbeat-evoked responses, i.e. transients of neural activity occurring in response to each heartbeat, during imagination. The amplitude of heartbeat-evoked responses differed between imagining oneself and imagining a friend, in the precuneus, mid and posterior cingulate regions bilaterally. Effect size was modulated by the general daydreaming frequency of participants but not by their interoceptive abilities. These results could not be accounted for by other characteristics of imagination (e.g., the ability to adopt the perspective, valence or arousal), nor by cardiac parameters (e.g., heart rate) or arousal levels. Heartbeat-evoked responses thus appear as a neural marker distinguishing self from other during imagination.

**Highlights:** - Heartbeat-evoked responses differentiate self from other during imagination.

- These effects were located in the precuneus and mid- to posterior cingulate.

- The neural monitoring of the body could be a mechanism for self/other distinction.

## Introduction

Experimentally, heartbeat-evoked responses (HERs), i.e. transient changes in brain activity locked to heartbeats, were shown to encode the degree of self-relatedness of spontaneous thoughts (Babo-Rebelo et al., 2016a, 2016b) and were associated with different aspects and levels of bodily self-consciousness (Park et al., 2016; Sel et al., 2016). These results suggest a link between the neural monitoring of the heart and selfhood, such that the brain would refer to internal bodily signals, in particular the heart, to tag mental processes as being self-related. However, previous experiments explored *degrees* of selfhood or used paradigms involving external sensory stimuli. If HERs contribute to a general mechanism for the implementation of the self, they should also distinguish between self and other, during internally-generated imagination. Interestingly, some of the regions that distinguish self from other in perspective taking (Vogeley and Fink, 2003) and in particular during imagination (Ruby and Decety, 2001), i.e. the precuneus and anterior insula, also respond to heartbeats (Babo-Rebelo et al., 2016a, 2016b; Park et al., 2017). Here, we test the hypothesis that heartbeat-evoked responses distinguish between self and other during imagination.

To test this hypothesis, we recorded both brain activity using magnetoencephalography (MEG) and cardiac activity (electro-cardiogram, ECG) while participants had to imagine themselves or a friend, in a series of cued scenarios (Fig.1A). Participants imagined themselves from the first-person perspective, from inside their body, and imagined their friend from the third-person perspective, by picturing the friend in the scenario. The scenarios either explicitly cued an action (ex: “petting a tiger”), or indicated a general context (ex: “in a space rocket”). All scenarios were unlikely or unreal, to avoid between-condition differences in memory retrieval and familiarity. After imagining each scenario, participants were asked to rate in 5-point scales how well they were able to adopt the indicated perspective (Perspective scale), the valence of the imagined scenario (Valence scale) and their level of arousal during imagination (Arousal scale). We tested whether the amplitude of heartbeat-evoked responses (HERs) during imagination of oneself differed from the amplitude of HERs taking place during imagination of the friend (Fig.1B).

**Figure 1:**
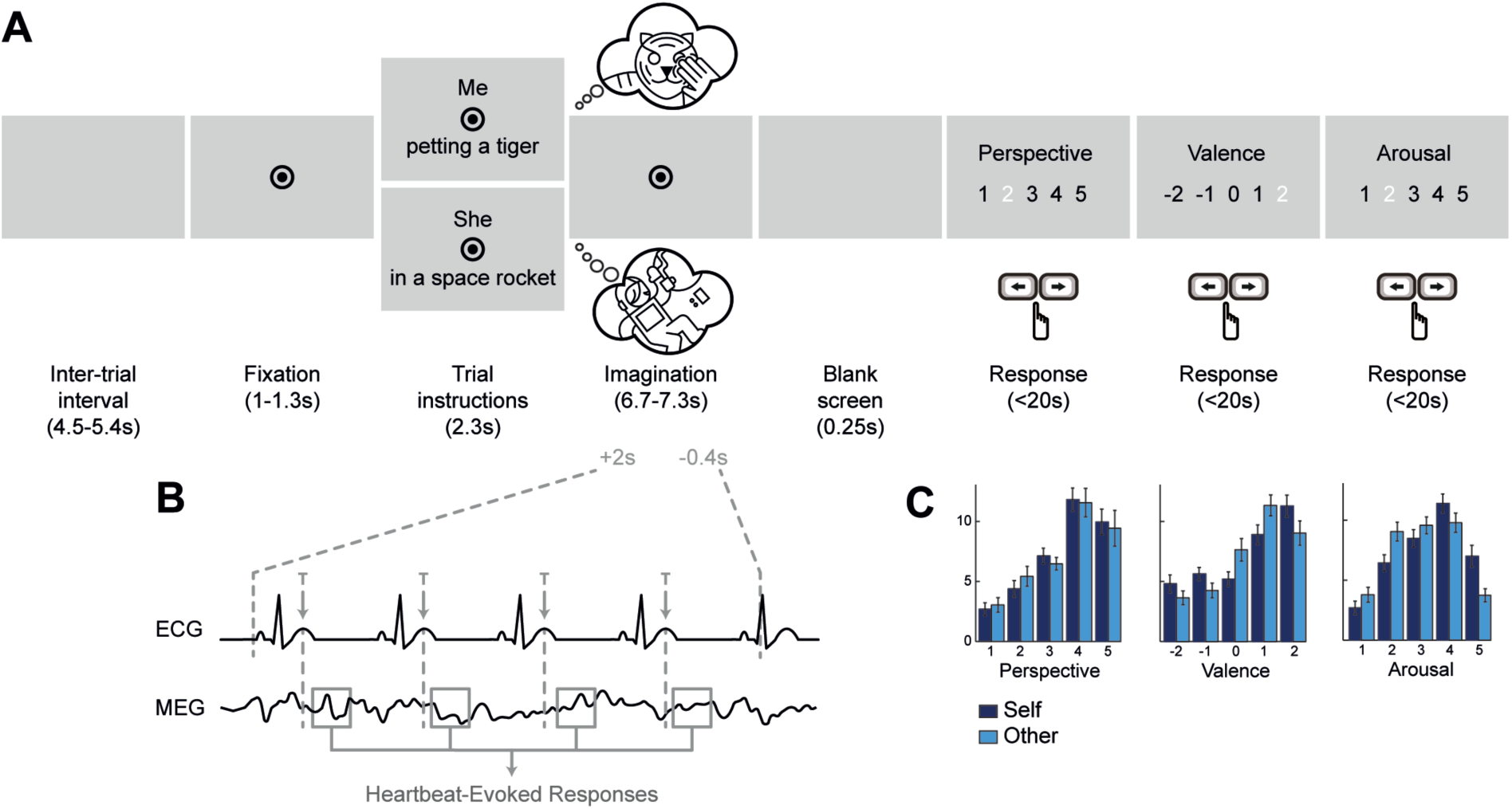
Experimental paradigm and behavior. ***A***, Time course of a trial. At each trial, participants had to imagine the person (Self, i.e. oneself from the first-person perspective, or Other, i.e. a friend from the third-person perspective) in the scenario indicated, until fixation disappeared. They then had to rate the imagined scenario in terms of Perspective (how well they succeeded in adopting the indicated perspective), Valence (how pleasant the scene was) and Arousal (how arousing the scene was). ***B***, Computation of Heartbeat-Evoked Responses (HERs) during the imagination period. T-peaks occurring from 2s after the beginning of the imagination period to 0.4s before the end of this period were selected. MEG data was extracted locked to these T-peaks to compute HERs. ***C***, Distribution of responses for the Perspective, Valence and Arousal scales, for both Self and Other trials, across all participants. Self trials were significantly more arousing than Other trials (paired t-test on the average Arousal ratings for Self and Other: p=0.0005, uncorrected). Error bars indicate SEM.

## Materials & Methods

### Participants

25 right-handed volunteers participated in this study after giving written informed consent. They were paid for their participation. The study was approved by the local ethics committee. Participants were screened to exclude cases of prosopagnosia or any cardiac problems. All participants were native French speakers. Two participants were excluded from analysis, one because of a noisy electrocardiogram recording, and the other because of an extremely fast heart-rate (mean interbeat interval of 555ms, >2 SDs faster than the average interbeat interval in the other participants). Twenty-three participants were thus included in the analysis (9 male; mean age: 24.3±0.6).

### Experimental procedure

The day before the experiment, participants were asked to choose the friend they would imagine in the task. The friend had to be the same gender and around the same age as the participant. The participant had to know him/her quite well and had to be able to clearly visualize him/her. It could not be someone the participant was romantically involved with, their best friend nor a relative. To assess the closeness of the selected friend, participants filled in a modified version of the Relationship Closeness Inventory (Berscheid et al., 1989) (RCI, excluding questions related to romantic relationships), where total scores range from 3 to 30. The average RCI score among participants was 12.4±0.8, which is intermediate between close (scores around 16) and distant relationships (scores around 9).

Before the MEG recording, participants were given written and oral instructions about the task, described in the next section. They performed a short practice block (2 trials of each condition), followed by four blocks of 9 trials of each condition (randomly presented), during which MEG and physiological data were acquired. Then, participants performed the heartbeat-counting task, in order to assess their interoceptive abilities (Schandry, 1981). They had to count their heartbeats while fixating the screen, during one practice block followed by five test blocks of different durations (practice block: 45s; test blocks: 60, 80, 100, 120, 140s – order randomized between participants), without feedback on performance. The heartbeat perception score was computed as the mean score over the 5 test blocks. In the same session, participants also performed a trait-judgment task and a resting state recording. These data are not presented here. After the recording session, participants completed a short feedback questionnaire and the Daydreaming Frequency Scale (Giambra, 1993; Stawarczyk et al., 2012).

### Imagination task

Each trial (Fig. 1A) began with a fixation mark (central black dot, radius 0.21° of visual angle, surrounded by a black circle, radius 0.52° of visual angle), presented for 1 to 1.3s on a gray background, followed by the instruction screen. The instruction screen specified the person to imagine (condition Self: “Me”, or condition Other: “He”/”She”), above fixation, and provided a brief description of the scenario to be imagined, below fixation. The instructions remained on screen for 2.3s, after which they disappeared, leaving only the fixation mark. Participants had then to imagine the scenario while fixating, until the fixation mark disappeared, after 6.7 to 7.3s.

Participants were instructed to adopt a first-person perspective in trials where they had to imagine themselves, meaning they should imagine the scenario from inside their own body. In trials where they had to imagine their friend, participants should adopt a third-person perspective and visualize the friend without interacting with him/her. Participants were also instructed to focus on the imagination of the person rather than on the visual details of the scenario.

The imagination period stopped with a blank screen (0.25s) and was followed by the presentation of the three scales (the order was randomized between participants, but constant for each participant). Participants had to rate on 5-point scales: the perspective (“How well did you manage to adopt the perspective in this scenario?”, from 1: not very well, to 5: very well); the valence (“How pleasant was the scenario?”, from −2: very unpleasant, to 2: very pleasant); and the arousal (“How arousing was the imagined scenario?”, from 1: not arousing, to 5: very arousing) of the imagined scenario. The order of presentation of the responses (ascending or descending) was randomized at each trial, to minimize motor preparation. Participants responded by pressing left and right buttons (index and middle finger respectively) to select the appropriate response. They validated their response with their right thumb, within 20s per scale. A new trial started after an inter trial interval (blank screen, 4.5 to 5.4s). The task was displayed on a semi-translucent screen at 85 cm viewing distance and was programmed using PsychToolbox.

### Scenarios

A list of 72 scenarios, e.g. a brief description of the situation to be imagined, was created, 42 of which contained an action verb (examples: “to drive a Formula 1 car”, “to erect a standing stone”). The remaining scenarios did not contain any verb, and rather indicated a context or environment (examples: “in the Middle Ages”, “in the jungle”). Scenarios described unreal or unlikely situations, which participants most likely never lived before. Each scenario was presented only once during the experiment. Scenarios assigned to condition Self for subject 1 were assigned to condition Other for subject 2 and vice-versa, for all pairs of subjects. This way, each scenario was associated with “Self” and with “Other” conditions the same number of times across subjects. The proportion of scenarios with and without a verb was distributed equally between conditions.

### Recordings

Continuous magnetoencephalographic (MEG) data were acquired using a whole-head MEG system with 102 magnetometers and 204 planar gradiometers (Elekta Neuromag TRIUX, sampling rate of 1000Hz, online low-pass filtered at 330Hz). Electrocardiogram data (ECG, 0.03-330Hz) were obtained from 7 electrodes placed around the base of the neck and referenced to a left abdominal location. The ground electrode was located on the back of the neck. Two ECG electrodes were placed over the left and right clavicles, two over the top of the left and right shoulders, two over the left and right supraspinatus muscle and one over the upper part of the sternum. The electrodermal activity (EDA, two electrodes on the sole of the left foot) as well as respiratory activity (respiratory belt positioned around the chest, at the level of armpits, respiratory transducer TSD201 BIOPAC system) were also recorded (low-pass filtered at 330Hz). Electromyographic activity (EMG, two electrodes on the right cheek, 10-330Hz) from the right zygomaticus major was acquired in order to control for facial muscle activity, a potential source of noise for MEG data. Indeed, because the scenarios were quite unlikely, participants might occasionally smile or laugh. Horizontal and vertical eye position and pupil diameter were monitored using an eye-tracker (EyeLink 1000, SR research) and recorded together with electrophysiological data.

### MEG data preprocessing

Continuous MEG data were denoised using temporal signal space separation (TSSS, as implemented in MaxFilter) and filtered between 0.5 and 45Hz (4^th^ order Butterworth filter). Large movement or muscle artefacts were visually detected and the corresponding data excluded from analysis. Independent Component Analysis (ICA) as implemented in the Fieldtrip toolbox (version: 20161025) (Oostenveld et al., 2011) was used to correct for the cardiac field artifact, on both magnetometers and gradiometers, based on epochs of −0.2 to 0.2s around the R-peaks of interest devoid of movement, muscle, blink or saccade artefacts. Because TSSS induces rank-deficiency, we defined the number of ICA components by first computing a Principal Component Analysis (PCA). We then removed all independent components with mean pairwise phase consistency (Vinck et al., 2010) with the ECG in the 0-25 Hz range larger than two standard deviations of all components. We iterated this procedure until no outlier components were found or a maximum of two excluded components was reached. A similar ICA procedure was then applied to correct for blink artefacts. Each trial was divided in five segments (from the onset of fixation to the end of the imagination period), devoid of large saccades (>4°), movement or muscle artefacts. Component decomposition was performed on those segments, with the number of components defined by PCA. We then computed the correlation between the time course of each component and the vertical EOG signal. Using an iterative procedure, we rejected components whose correlation coefficient exceeded three standard deviations of all components, until no outlier was present or the maximum of three components to reject was reached. For two subjects the automatic selection of components was ineffective and one subject did not have a vertical EOG recording. For these subjects, the selection of components was done by visually identifying the characteristic topographies of blink ICA components and rejecting those components.

### Heartbeat-evoked responses (HERs)

R- and T-peaks were detected on derivation lead II of the ECG, except in two subjects for whom T-peaks were not clearly visible on lead II and therefore were detected on lead III. ECG recording was band-pass filtered, between 0.5 and 40 Hz (4th order Butterworth filter). We first detected the R-peaks, by correlating the ECG with a template QRS complex defined on a subject-by-subject basis and by identifying the local maximum within the episodes of correlation larger than 0.7. T-peaks were then detected by first, correlating the ECG with a template of the T-peak; second, identifying the local maxima within episodes of correlations above a certain correlation value (adapted for each subject) that followed an R-peak by at most 0.4s. R- and T-peak detection was visually verified in all subjects. R- and T-peaks corresponding to noisy ECG data (movement artefacts) or followed/preceded by extra-systolic events were excluded from analysis.

The T-peaks occurring during the imagination period (from 2 seconds after the beginning of the period, to −0.4s before the end of the imagination period) were used for HER analysis. Epochs (from 0.1s before to 0.4s after the selected T-peaks) contaminated with movement or muscular (in particular of the zygomaticus, recorded with the EMG) artifacts were not included in the analysis. peaks which were followed by an R-peak by less than 0.4s were not considered for HER computation to avoid any overlap between the HER window of analysis and the residual R-peak artefact. Artefact-free HERs corresponding to Self and Other trials were computed by averaging across heartbeats magnetometer data low-pass filtered at 30 Hz (4^th^ order Butterworth filter). Only trials for which a response was recorded for each of the three scales were included in the analysis. The time window retained for analysis was 80 to 350 ms after the T-peak, during cardiac relaxation when the cardiac artefact is minimal (Dirlich et al., 1997).

### Statistical analyses

The difference in HERs between Self and Other was tested on magnetometer data, in the artifact-free time window 80-350ms after the T-peak, using a cluster-based permutation t-test (Maris and Oostenveld, 2007). This method does not require any a priori on spatial regions or latencies and intrinsically corrects for multiple comparisons in time and space.

Briefly, the procedure entails the following processing steps. A paired *t-*test was performed to compare HERs for Self versus Other. Individual samples whose *t-*value was below a threshold (p<0.05, two-tailed) were clustered together based on temporal and spatial adjacency (with a minimum of 4 neighboring channels). A candidate cluster was characterized by the sum of the *t*-values across the individual samples. To test whether such a cluster could be obtained by chance, we permuted the labels “Self” and “Other” 10,000 times and selected the maximal positive cluster-level statistic and the minimal negative cluster-level statistic at each randomization. The two-tailed Monte-Carlo *p*-value corresponds to the proportion of elements in the distribution of maximal (or minimal) cluster-level statistics that exceeds (or is inferior to) the originally observed cluster-level test statistics. The amplitude of the cluster corresponds to the average of magnetometer data across the sensors and time window showing a significant difference between conditions.

### General linear models

General linear models (GLM) were computed to better characterize the trial by trial HER cluster amplitude. Importantly, GLMs assess the unique contribution of each regressor, since the shared variance between regressors is partialled out. Trial by trial HER cluster amplitude and regressors (ratings on the scales Perspective, Valence and Arousal) were z-scored prior to GLM computation. The regressor “condition” was coded with the value 1 for Self trials, and the value −1 for Other trials. An additional regressor corresponding to the trial number was introduced, in order to group HERs belonging to the same trial. GLMs were computed for each subject, and the betas corresponding to each regressor of interest were tested against zero, over subjects.

### Bayes factors to assess evidence in favor of the null hypothesis

Bayes factors were computed to evaluate evidence in favor of the null hypothesis. We computed the maximum log-likelihood of the model in favor of the “null” hypothesis and the model in favor of the “effect” hypothesis. The group-level random-effect Bayes factor was computed with the prior reference effect corresponding to an effect differing from 0 under a *t*-test with a *p* value of 0.05. We then used the Bayesian information criterion to compare the two models and compute the corresponding Bayes factor. Bayes Factor interpretation was done according to (Kass and Raftery, 1995). Bayes factors for correlation were computed using an online calculator tool (http://pcl.missouri.edu/bayesfactor, with the version 0.9.8 of the BayesFactor package, R version 3.3.2 (2016-10-31) on i386-redhat-linus-gnu).

### Surrogate heartbeats

To test whether the observed effects were truly locked to heartbeats, we checked whether differences between Self and Other trials could be obtained with a sampling of neural data that was desynchronized from heartbeats. We created 1,000 permutations of heartbeats, where the timings of the heartbeats of trial *i* in the original data were randomly assigned to trial *j*. The same analysis was performed, with the same criteria for rejecting artefactual epochs and computing HERs. For each permutation, we obtained a set of neural responses to surrogate heartbeats and computed the cluster summed t-statistics as described above. For each permutation we extracted the smallest negative sum of t-values, and compared the distribution of those surrogate values with the observed original (negative) sum of t-values.

### Anatomical MR acquisition and preprocessing

An anatomical T1 scan was acquired for 22 out of 23 participants. Segmentation of the data was processed with automated algorithms provided in the FreeSurfer software package (Fischl et al. 2004) (http://surfer.nmr.mgh.harvard.edu/). Segmentations were visually inspected and edited when necessary. The white-matter boundary was determined using FreeSurfer and was used for subsequent minimum-norm estimation.

### Source reconstruction

We reconstructed sources of HERs occurring from 2 to 4s after the onset of the imagination period. Source reconstruction and surface visualization were performed with the BrainStorm toolbox (version 13-Dec-2016, on Matlab R2012b) (Tadel et al., 2011). For the participant who did not have an anatomical scan, we warped the ICBM152 anatomical template (http://bic.mni.mcgill.ca/ServicesAtlases/ICBM152NLin2009) to fit the shape defined by the digitized head points obtained before MEG acquisition. After co-registration between the individual anatomy and MEG sensors, cortical currents were estimated using a distributed model consisting of 15,002 current dipoles from the combined time series of magnetometer and gradiometer signals using a linear inverse estimator (weighted minimum-norm current estimate, signal-to-noise ratio of 3, Whitening PCA, depth weighting of 0.5) in an overlapping-spheres head model. Dipole orientations were constrained to the individual MRIs. Cortical currents were then averaged over the HER time windows for which a significant difference between Self and Other was identified in sensor space, spatially smoothed (FWHM 7mm) and projected to a standard brain model (ICBM152, 15,002 vertices).

Reliable differences in dipole current values were identified using the cluster-based procedure (first-level p-value: 0.01, 1 neighboring vertex, 1,000 randomizations) applied to the 15,002 vertices, as described for the sensor level analysis. The obtained Monte-Carlo p-value was thus intrinsically corrected for multiple comparisons across vertices.

Analyses at the source level in a given region of interest were carried by extracting the data of each vertex and flipping the sign of vertices that differed from the main sign of the region. This was achieved using Brainstorm functions for sign flipping. To select the vertices corresponding to the insula, we used the masks from (Deen et al., 2011) corresponding to the ventral anterior, dorsal anterior and posterior insula bilaterally and transformed them to fit the anatomical template of Brainstorm, using the function ImCalc of SPM12. Differences between Self and Other over time, for each insular region of interest, were assessed by computing a cluster-based permutation test, over time (first-level p-value: 0.05; number of randomizations: 1,000). This method generated candidate clusters, for which we computed Cohen’s d effect sizes.

### Cardiac measures

Interbeat intervals (IBIs) consisted of the average time distance between T-peaks in the imagination period. The heart rate variability (HRV) corresponded to the standard deviation of the interbeat intervals.

### Arousal controls

We used two different measures of arousal: pupil diameter, and the arousal ratings given by participants at each trial.

Pupil diameter was obtained with the Eyelink software (EyeLink 1000, SR research). Blink epochs were automatically identified by the acquisition software. We extended those epochs by 80ms before and after to ensure that the whole blink event was included. We also identified noise in pupil diameter data (e.g. signal variation larger than 1 in arbitrary units, in a 300ms time window). Portions of pupil diameter data containing blinks or noise were linearly interpolated and a low-pass filtered at 10Hz (fourth-order Butterworth filter). Pupil data were then epoched from 2 to 4 seconds after the onset of the imagination period. Epochs with more than 30% of interpolated data were excluded from further analysis. The remaining epochs were z-scored. Four subjects were excluded for having a too low number of remaining trials (< 1.5 SD).

Trials in the Self condition were rated as more arousing than trials in the Other condition. To verify that the HER difference was not due to a difference in perceived arousal, we employed a stratification procedure which consists in removing trials until we obtain two groups of trials in the Self and Other condition with similar arousal ratings. The stratification procedure ran as follows. In each subject, the mean arousal rating was computed for Self and Other. One of the two conditions was then randomly selected. If the mean rating of the selected condition was larger than the mean rating of the non selected condition, the trial corresponding to the largest arousal rating of the selected condition (or a trial randomly chosen among all trials having the maximum arousal rating) was excluded. Conversely, if the mean arousal rating of the selected condition was smaller, the trial corresponding to the smallest arousal rating (or a trial randomly chosen among all trials having the minimum rating) was excluded. This procedure was iterated until the difference in arousal ratings between conditions was smaller than 0.02. Overall, 61.35 (±1.47) out of 72 trials were included (minimum number of trials included: n=46; maximal: n=69), and no difference was found between conditions in the final number of trials included (paired t-test: t_(22)_=0.066, p=0.95, Bayes Factor = 4.12 – substantial evidence for H0). We then tested whether cluster amplitudes and pupil diameter still differed between conditions in this subset of trials that did not differ in terms of arousal ratings

## Results

### Behavioral results

The distributions of ratings on the Perspective, Valence and Arousal scales for the Self and Other conditions are presented in Figure 1C. Mean ratings did not differ between Self and Other conditions in the Perspective or Valence scales (Perspective: mean Self: 3.6±0.1 SEM, mean Other: 3.5±0.1, paired t-test Self x Other, t_(22)_=0.7, p=0.5, Bayes Factor (BF) = 2.80 – anecdotal evidence for_1_ H0; Valence: Self: 3.5±0.1, Other: 3.5±0.1, t_(22)_=−0.5, p=0.6, BF=3.36 – substantial evidence for H0; uncorrected p-values). Imagining oneself was rated as being significantly more arousing than imagining the other (Arousal: Self: 3.4±0.1, Other: 3±0.1, t_(22)_=4.10, p=0.0005, BF=549.3 – decisive evidence for H1; uncorrected p-value).

### HER amplitude differs between self and other

We compared the amplitude of HERs occurring during imagination of self with the amplitude of HERs occurring during imagination of other, for heartbeats (T-peaks) occurring between 2s after the onset of the imagination, to −0.4 seconds before the end of the imagination period (Fig. 1B). We excluded the beginning of the imagination period to make sure participants already started imagining the scenario.

Our whole-brain, whole time-window analysis of HERs showed that HERs significantly differed over posterior sensors (Fig. 2A), in the time window 307-326ms after the T-peak (Fig. 2B; cluster sum(t)=−733.713, Monte-Carlo p=0.012).

**Figure 2:**
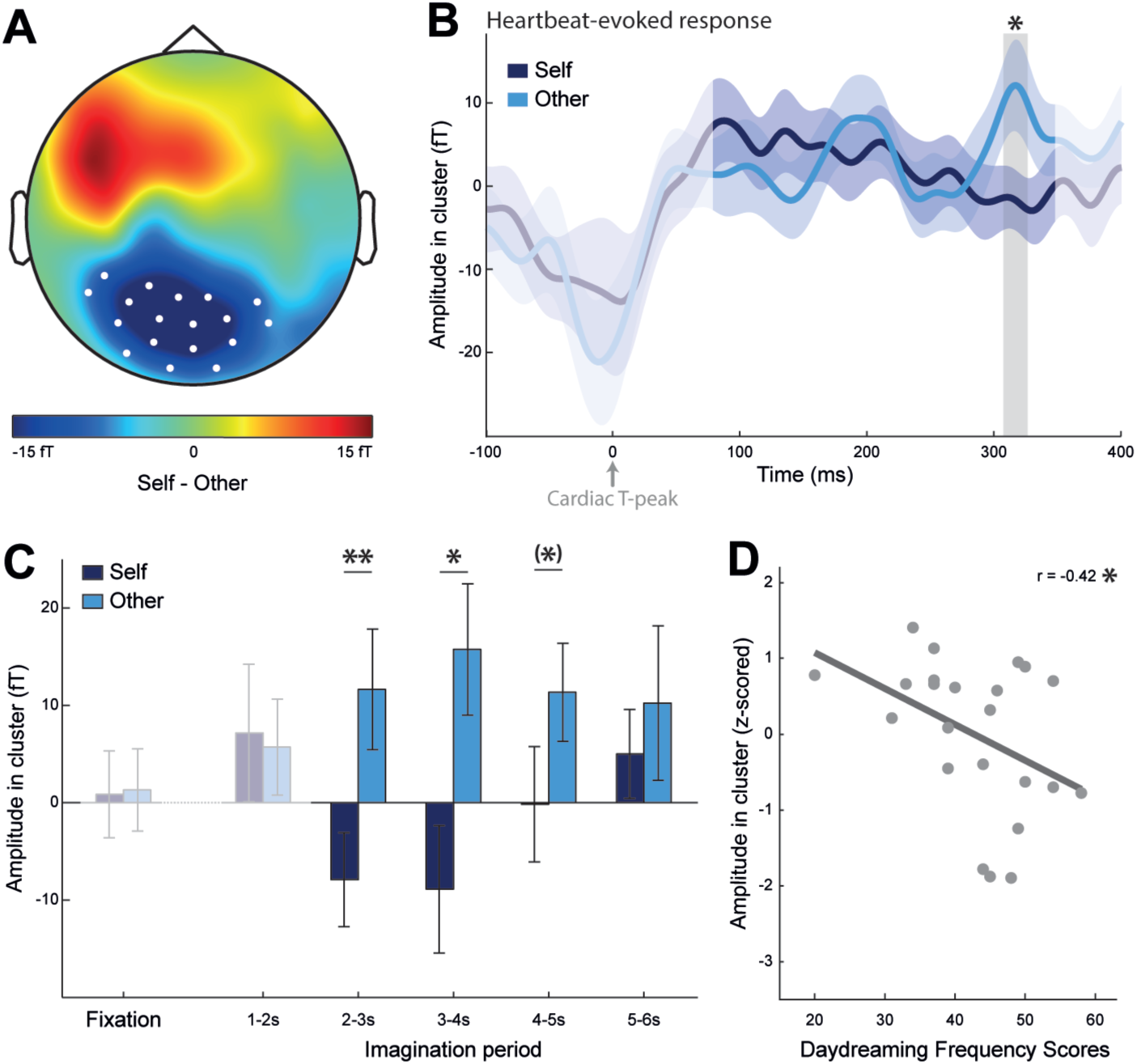
Differential Heartbeat-Evoked Responses (HERs) for imagining oneself (Self) or a friend (Other). ***A***, Topographical map of the HER difference between Self and Other conditions, grand-averaged across 23 participants, in the 307-326ms time window in which a significant difference was observed (Monte-Carlo p=0.012). White dots represent the sensors contributing to the significant cluster. ***B***, Time course of the HER (±SEM) for Self and Other, averaged over the sensors marked in white in A. The signal that might be residually contaminated by the cardiac artifact appears in lighter color and was not included in the epoch analyzed. The grey area represents the time window in which a significant difference was observed. ***C***, Temporal evolution of the HER effect, during the imagination period. Amplitude in cluster corresponds to the average brain activity in the T-peak locked time window and sensors revealing a significant HER effect (sensors indicated in A, time window indicated in B). Cluster amplitude was computed for HERs occurring during fixation (1-1.3s), and during the imagination period divided in five windows of 1 second (1-2s, 2-3s, 3-4s, 4-5s, 5-6s). The largest cluster amplitude differences between Self and Other were observed in the windows 2-3s and 3-4s. ***D***, Correlation between the size of the HER effect and Daydreaming Frequency Scores (p=0.049). HER effect size was computed as the difference between the HER cluster amplitude for Self minus the HER cluster amplitude for Other, z-scored, for HERs in 2-4s of imagination period. Each dot represents one participant. **: p<0.01, *: p<0.05, (*): p< 0.1

We then looked at the temporal evolution of the effect, during the imagination period, in time windows of 1 second (Fig. 2C). The difference between Self and Other was most pronounced for T-peaks occurring in the time window from 2 to 4 seconds after the onset of imagination. We thus retained this interval for further analysis, although a two-way repeated measures ANOVA on the cluster amplitude, with the factors Condition (Self, Other) and Timing (2-3, 3-4, 4-5, 5-6 seconds) as factors, did not reveal a significant interaction between Condition and Timing (main effect of Condition: F_(1,22)_=11.81, p=0.0024; main effect of Timing: F_(3,66)_=0.33, p=0.80; interaction: F_(3,66)_=1.41, p=0.25).

Because the scenarios could indicate a context or directly cue an action through a verb, we verified that the effect was present in both types of scenarios (paired t-test between Self and Other, for context scenarios: t_(22)_=−2.31, p=0.031; for action scenarios: t_(22)_=−3.08, p=0.0055). In addition, there was no main effect of the type of scenario nor any interaction between the scenario type and the condition (two-way repeated measures ANOVA on the cluster amplitude, with factors Condition (Self, Other) and Scenario type (context, action): main effect of Condition: F_(1,22)_=11.25, p=0.0029; main effect of Scenario Type: F_(1,22)_=2.62, p=0.12; interaction: F_(1,22)_=0.40, p=0.53).

We next tested whether the HER difference between Self and Other was related to the ratings participants provided on a trial-by-trial basis on the success in adopting the perspective, the valence and the arousal of the imagined scenario. To test the unique contribution of the condition (Self/Other), the scales and a possible interaction between them, we performed a general linear model (GLM) for each subject, where HER cluster amplitudes were predicted by 8 regressors: the condition, ratings on each of the three scales, interaction between condition and each scale ratings and trial number. Only the betas corresponding to the regressor “condition” were significantly different from zero across subjects (t-test against zero for each β value corresponding to each regressor: Condition: β=−0.21, t_(22)_=−2.72, p=0.013, BF=12.9 – strong evidence for H1; Perspective ratings: β=−0.047, t_(22)_=1.76, p=0.092, BF=1.68 – anecdotal evidence for H1; Valence ratings: β=0.017, t_(22)_=1.00, p=0.33, BF=2.02 – anecdotal evidence for H0; Arousal ratigns: β=0.013, t_(22)_=0.70, p=0.49, BF=2.88 – anecdotal evidence for H0; Condition*Perspective: β=0.027, t_(22)_=1.59, p=0.13, BF=1.22 – anecdotal evidence for H1; Condition*Valence: β=0.015, t_(22)_=0.92, p=0.37, BF=2.26 – anecdotal evidence for H0; Condition*Arousal: β=−0.0046, t_(22)_=− 0.27, p=0.79, BF=3.92 – substantial evidence for H0). This indicates that HER cluster amplitude variations are uniquely explained by the person who is being imagined.

### Control for arousal effects

Self trials were rated as being more arousing than Other trials. Although the GLM analysis showed that arousal ratings did not modulate the difference in HERs between Self and Other, we ran two additional controls.

We first tested pupil diameter, which is considered as a physiological marker of arousal (Murphy et al., 2011). In this experiment, pupil diameter significantly correlated with arousal ratings (t-test against zero for Fisher-z-transformed Pearson correlation coefficients between arousal ratings and pupil diameter averaged over 2-4 seconds across subjects: r_(17)_=0.18, t_(17)_=4.26, p=0.00047). However, in the time window 2-4 seconds, where the HER effect is stronger, we did not find conclusive evidence for a difference in pupil diameter between Self and Other (t_(18)_=1.65, p=0.12, BF=1.34 – anecdotal evidence for H1).

We then further tested arousal as rated by the subjects at each trial. We performed a stratification procedure where some trials were rejected so as to cancel out the difference in arousal ratings between Self and Other. Using this procedure, the significant difference in arousal ratings disappeared (paired t-test between average ratings for Self vs Other, before stratification: t_(22)_=4.09, p=0.0005; after stratification: t_(22)_=0.14, p=0.89, BF=4.09 – substantial evidence for H0), the trend for a pupil difference disappeared (paired t-test on pupil diameter between conditions before stratification: t_(22)_=1.65, p=0.12; after stratification: t_(22)_=1.04, p=0.31, BF=1.95 – anecdotal evidence for H0) while the difference in cluster amplitudes remained robust (cluster amplitude difference between conditions before stratification: t_(22)_=−3.25, p=0.0036; after stratification: t_(22)_=−2.99, p=0.0068, BF=24.94 – strong evidence for H1; uncorrected p-values). These results confirm that differential arousal levels between Self and Other trials cannot explain the HER effects.

### HER effects and inter-subject variability

We tested whether the amplitude of the HER difference between Self and Other correlated with participants’ tendency to daydream in their daily lives, as assessed with a questionnaire (Giambra, 1993; Stawarczyk et al., 2012) after the experiment. We found a negative correlation between the daydreaming scores and the HER effect size, measured as the difference between HER cluster amplitude for Self and HER cluster amplitude for Other (Fig. 2D; daydreaming frequency scores: 42.74±1.82; Pearson correlation with the z-scored effect size: r_(21)_=−0.42, r^2^=0.172, p=0.049). This means that people who are used to daydreaming more have a larger HER difference between self- and other-imagination.

We also tested whether the size of the HER effect correlated with individual interoceptive abilities, as measured with the heartbeat counting task (Schandry, 1981). We found no evidence that interoceptive abilities modulate the amplitude difference between HERs for Self and HERs for Other (heartbeat perception scores: 0.76±0.029; Pearson correlation with the z-scored effect size: r_(21)_=0.038, r^2^=0.001, p=0.86, BF=2.61 – anecdotal evidence for H0).

### HERs in the anterior precuneus and mid/posterior cingulate cortex are responsible for these effects

To identify the regions generating the differential HERs, we reconstructed HER sources for Self and Other, averaged the reconstructed neural currents in the time window where we found an effect (307-326ms after the T-peak) and performed a cluster-based permutation test over all 15,002 vertices to compare activations for Self and Other. The differential HER amplitude was generated in a bilateral region centered on the posteromedial cortex, more precisely in the anterior precuneus and extending ventrally to the mid and posterior cingulate cortex (Fig. 3A, 3B; Table 1; Left: cluster sum(t)=482.01, Monte-Carlo p=0.022, mean Cohen’s d=0.71±0.009, cluster surface 22.82cm^2^; Right: cluster sum(t)=−600.39, Monte-Carlo p=0.016, mean Cohen’s d=−0.74±0.008, cluster surface 26.27cm^2^).

**Figure 3:**
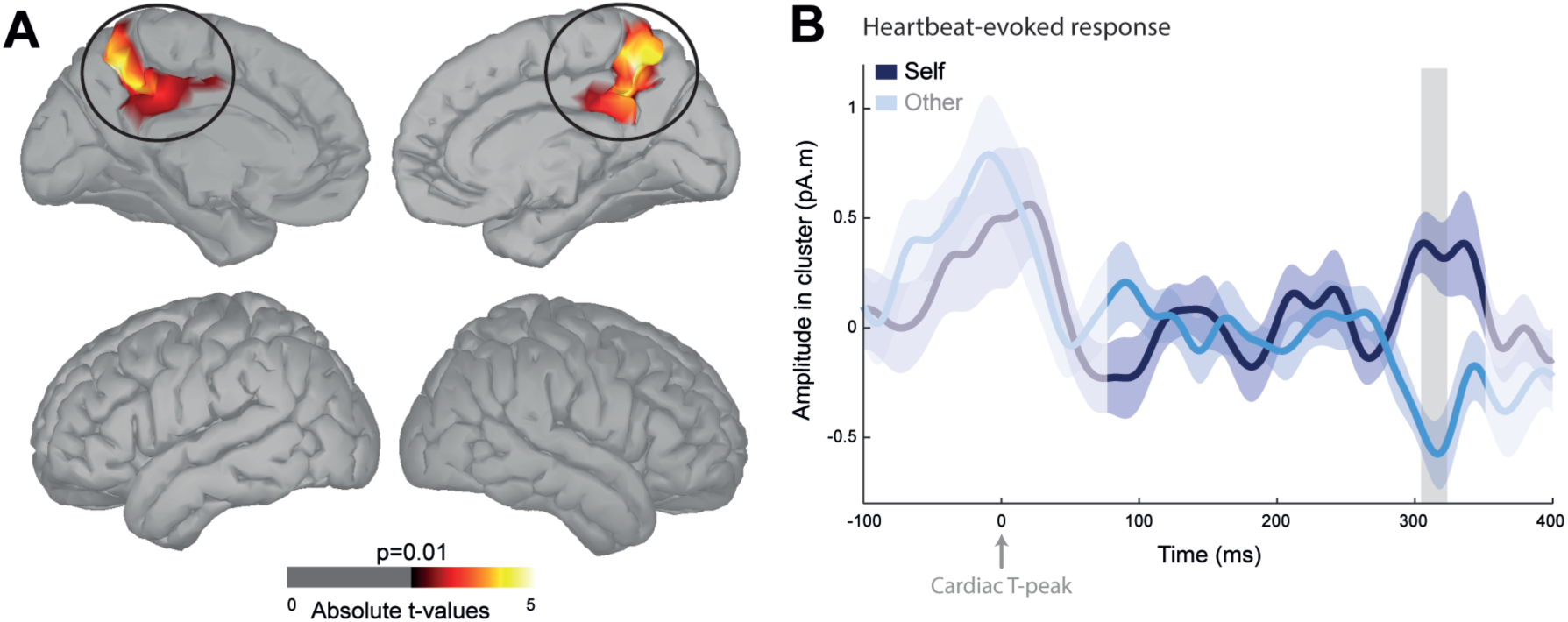
Neural sources of the Heartbeat-Evoked Response (HER) effects. ***A***, HER differences between Self and Other were localized in the anterior precuneus and mid/posterior cingulate cortex bilaterally (left: Monte-Carlo p=0.022; right: Monte-Carlo p=0.016; threshold for visualization: >30 contiguous vertices at uncorrected *p*<0.010). ***B***, Time course of the HERs (±SEM) in the region in A. The signal that might be residually contaminated by the cardiac artifact appears in lighter color. The grey area represents the time window in which a significant difference was observed at the sensor level.

**Table 1:** Anatomical description of the source clusters showing significant differential HERs (Fig. 3A).

| AAL regions | Peak *t* | Size (cm<sup>2</sup>) | MNI coordinates<br>(peak *t*) |  |  |
| --- | --- | --- | --- | --- | --- |
|  |  |  | X | Y | Z |
| Cluster left hemisphere |  |  |  |  |  |
| Precuneus | 4.99 | 10.12 | -10 | -58 | 50 |
| Mid-cingulate | 4.07 | 10.67 | -1 | -45 | 41 |
| Posterior cingulate | 2.96 | 1.20 | -1 | -44 | 29 |
| Cluster right hemisphere |  |  |  |  |  |
| Precuneus | -4.77 | 16.57 | 9 | -50 | 42 |
| Mid-cingulate | -4.02 | 7.63 | 16 | -47 | 35 |
| Posterior cingulate | -3.65 | 1.35 | 1 | -51 | 29 |
| Superior parietal gyrus | -2.87 | 0.066 | 14 | -54 | 61 |

### Region of interest analysis of the insula

The insular region did not come out as significant from the whole brain analysis. To probe this region further, we conducted a region of interest analysis of the insular cortex by extracting HER source data from three sub-regions of the insula (posterior, dorsal anterior and ventral anterior, bilaterally) as defined in (Deen et al., 2011), and tested for differences in HER amplitude between Self and Other over time, across the whole time window of interest (80-350ms relative to the T-peak). The clustering procedure identified some candidate clusters with a significant difference between Self and Other in the regions considered, but none of these clusters survived correction for multiple comparisons over time (Supplementary Figure; left: all |cluster sum(t)|<42, all Monte-Carlo p>0.25; right: all |cluster sum(t)|<62, all Monte-Carlo p>0.15; uncorrected p-values across multiple sub-regions tested).

We then compared the effect sizes within these candidate clusters with the effect sizes within the posteromedial region showing significant differences in HERs. Mean Cohen’s d across vertices in the left precuneus/cingulate cluster was 0.71±0.009 (ranging from 0.59 to 1.04), and for the right homologue was 0.74±0.008 (ranging from −0.59 to −0.99). In the left insula, the largest mean Cohen’s d among candidate temporal clusters was 0.51. In the right insula, the largest mean Cohen’s d among candidate temporal clusters was 0.57 (absolute values).

Thus, the insula region does not show any reliable difference in HERs. The few candidate temporal clusters show effect sizes that are 23 to 28% smaller than the effect sizes observed for the posteromedial region. This indicates that the insula is not generating HERs that reliably distinguish Self and Other.

### Test of cardiac parameters during self- vs other-imagination

To account for possible concomitant differences in cardiac parameters, we compared the mean interbeat-interval (IBI) and heart-rate variability (HRV) in the time window 2-4s of imagination between Self and Other conditions. We found no evidence for a difference in IBIs (IBI for Self: 853.93ms±24.81; IBI for Other: 853.41±24.48; t_(22)_=0.25, p=0.81, BF=3.96 – substantial evidence for H0), and only a trend in HRV (HRV for Self: 48.98±2.54; HRV for Other: 53.14±3.75; t_(22)_= −1.82, p=0.082, BF=1.87 – anecdotal evidence for H1).

### These effects are time-locked to heartbeats and of neural origin

To show that this effect was truly locked to heartbeats and not driven by slow fluctuations of neural activity differing between conditions, we permuted heartbeat timings between trials 1,000 times and performed the same analyses on these surrogate heartbeats at the sensor level. Only 4 out of 1,000 permutations led to a cluster *t* statistic larger (in absolute values) than the original one, which indicates that our effect is an *evoked-response* to heartbeats, with a Monte-Carlo p value of 0.004.

Heart contractions generate electrical currents that create a magnetic field, which is directly picked up by MEG sensors. In order to show that our results are not due to the electrical activity of the heart but to brain activity, we compared the electrical activity of the heart as measured with the ECG between Self and Other trials, in the time window were we find the HER results. We did this for each of the seven vertical and horizontal ECG leads acquired, to best estimate possible heart-to-MEG signal propagations. We could not find any evidence for a significant difference in heart electrical activity between conditions (paired t-test between mean Self vs Other ECG amplitude averaged over 307-326ms relative to the T-peak in 2-4s of the imagination period: all |t_(22)_|<1.79, all uncorrected p>0.086; all Bayes Factors ≥ 1.15, anecdotal (4 tests) or substantial (10 tests) evidence for H0).

## Discussion

We hypothesized that the distinction between self and other during imagination could involve an internal mechanism based on the neural monitoring of heartbeats. Our results confirmed this hypothesis, by showing that the amplitude of heartbeat-evoked responses (HERs) in the anterior precuneus, mid and posterior cingulate cortices bilaterally differed between imagination of self and imagination of a friend. We also observed that participants who daydream more in their daily lives had larger HER amplitude differences between self and other. Overall, these results further support the proposal that the neural monitoring of internal signals could constitute a mechanism to tag mental processes as being self-related or not (Babo-Rebelo et al., 2016a, 2016b; Park and Tallon-Baudry, 2014; Tallon-Baudry et al., 2017).

### Specificity and neural origin of the HER difference between self and other

We verified that the difference in HER between self and other was uniquely explained by the person being imagined (oneself or the friend), and was not modulated by other factors (success in adopting the perspective, valence or arousal). Besides, the HER difference was equally found for scenarios involving an action and scenarios which did not involve an action. This suggests that this mechanism is not content specific, but could potentially account for any type of content of imagination, where a self vs. other distinction is to be made.

Because the electrical activity of the heart can be directly picked up by MEG sensors, we controlled that our results were of neural and not of cardiac origin. We also showed by a permutation procedure that our effects were truly locked to cardiac T-peaks and not due to sustained fluctuations of brain activity unrelated to heartbeats. Furthermore, we could not measure any changes in the cardiac activity itself. The results we obtained are thus due to fluctuations in the amplitude of a *neural* response to an internal stimulus, the heartbeat, which in itself does not seem to carry any relevant information other than its simple occurrence.

### The HER difference between self and other is not related to classical markers of explicit interoception

The HER is a measure of automatic, implicit interoception. We tried to relate the difference in HER between self and other to two markers of explicit interoception, i.e. interoceptive accuracy (Garfinkel et al., 2015; Schandry, 1981) and insular cortex activation (Critchley et al., 2004). There was no correlation between the size of the HER effect and individual interoceptive abilities as measured with the heartbeat counting task. Note that recent evidence suggests that this task, that has been widely employed, might be confounded by differences in heart rate between participants (Zamariola et al., 2018).

The region of interest analysis of the insular cortex did not reveal any differences in HER amplitude between self-and other-imagination. HER effects were previously found in the insula in the context of explicit interoceptive tasks, where participants were asked for instance to monitor their heartbeats (Canales-Johnson et al., 2015; Pollatos et al., 2005). A marginal effect was also found in relation to the self-relatedness of spontaneous thoughts (Babo-Rebelo et al., 2016b). Influential theories about the self have postulated that the insular cortex-in particular the anterior portion - would integrate interoceptive inputs with cognitive processing and thereby generate the sense of self (Craig, 2009). Here, we find no evidence supporting this proposal.

### The posteromedial cortex generates the HER difference between self and other

The region most reliably contributing to the HER difference between self and other in this task was the posteromedial cortex. The most ventral portion of this region, encompassing the posterior cingulate cortex, within the default network (Andrews-Hanna et al., 2014), has previously been shown to elicit HERs in relation to the self-relatedness of spontaneous thoughts (Babo-Rebelo et al., 2016a, 2016b) and bodily self-consciousness (Park et al., 2016). Evidence is thus now converging to support the idea of an integrative role of the posterior cingulate cortex, between self-related cognitive processes (Fox et al., 2015; Qin and Northoff, 2011; Vogt and Laureys, 2005) and physiological functions such as cardiac monitoring. This region is not a direct target of visceral signals and the pathways susceptible to conveying visceral information remain unknown. However, this structure is functionally connected to the insula (Zhang et al., 2014), and both functionally (Bzdok et al., 2015) and structurally (Parvizi et al., 2006) connected to the anterior cingulate cortex, two direct targets of visceral inputs (Critchley and Harrison, 2013).

The posterior cingulate cortex appears to be the core region for self-related HERs across studies, but we here also find self-related HERs in a more dorsal and anterior part of the precuneus. This effect might be associated with specificities of this task, in particular with the process of actively adopting a certain visual perspective (Zaehle et al., 2007).

### Self, other and perspective taking

Here, the two protagonists of imagined scenarios were associated with two distinct imagery perspectives: the first-person perspective - the self was imagined from within the body, and the third-person perspective - the friend was visualized in the scenario. We chose not to ask participants to imagine the friend from the first-person perspective, because that would be very unnatural, nor the self from the third-person perspective, because that would target a more distantiated self (Kross et al., 2005) thereby reducing the contrast between self and other (Christian et al., 2015). Self and other are indeed preferentially and intrinsically associated with a certain perspective.

However here, HERs were obtained not only for self but also for the other condition, with opposite signs indicating that they originate from different neural populations. Those previous experiments contrasted strong engagement against weak engagement of the self in thoughts. Here, we show that HERs do not simply detect the presence of an active protagonist, they signal *who* this active protagonist is, oneself or someone else.

## Conclusion

To conclude, our results suggest that HERs could be a mechanism to disentangle self from other, during an entirely internal mental process such as imagination. In externally-driven processes, sensory signals or motor feedback can characterize the self when we hear our own voice (Heinks-Maldonado et al., 2005) or perform a voluntary movement (Blakemore et al., 1999), for example. For internally-driven processes, heartbeats would work as a signal integrated by the brain to tag a mental process as being related to the self or not. The heartbeat itself does not carry any self-related information; the key process is the integration between the heartbeat input and ongoing brain activity. The precise mechanisms behind this integration are still unknown and will require further investigation to be elucidated.

---

**Supplementary figure:**
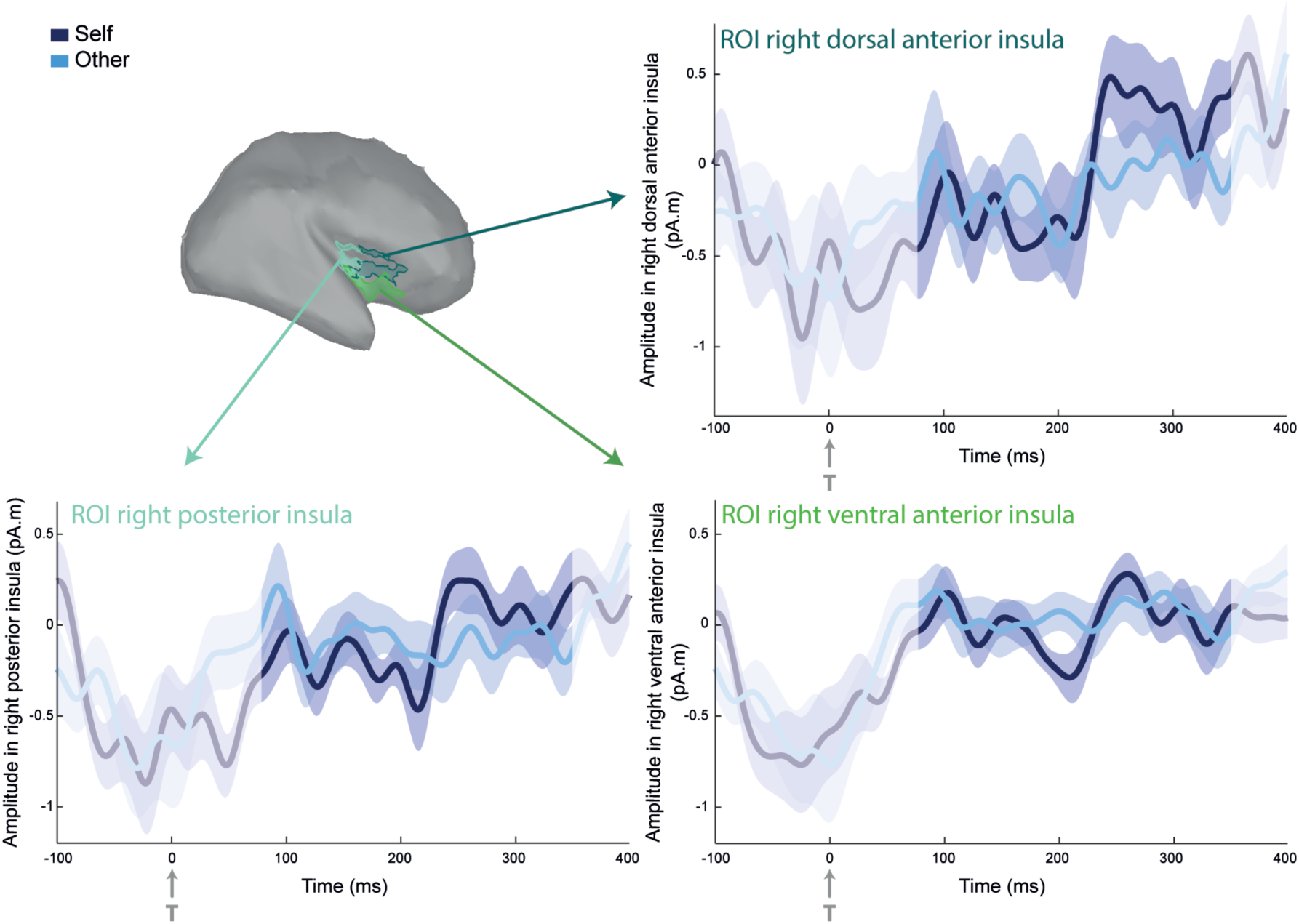
Heartbeat-Evoked Response (HER) in the three sub-regions of the right insular cortex. No significant differences were observed between HERs during self-vs other-imagination.

## Aknowledgements

The authors thank Margaux Romand-Monnier and Christophe Gitton for their help with data acquisition and Leonor Babo for the illustrations of figure 1.

Contributions
M.B-R. and C.T-B. designed the study. M.B-R. acquired the data. M.B-R. and C.T-B. analyzed the data, with the help of A.B. M.B-R. and C.T-B. wrote the paper

